# Comparison of simple RNA extraction methods for molecular diagnosis of hepatitis C virus in plasma

**DOI:** 10.1101/2022.05.17.492343

**Authors:** Sayamon Hongjaisee, Yosita Jabjainai, Suthasinee Sakset, Kanya Preechasuth, Nicole Ngo-Giang-Huong, Woottichai Khamduang

## Abstract

We evaluated the efficacy of four simple RNA extraction methods for the detection of hepatitis C virus (HCV) in plasma samples: silica-membrane-based, magnetic beads-based, boiling with Diethyl Pyrocarbonate (DEPC)-treated distilled water or a commercial lysis buffer. HCV RNA was detected using both real time reverse transcription polymerase chain reaction (RT-PCR) and reverse transcription loop mediated isothermal amplification (RT-LAMP). The magnetic beads-based extraction can be used as an alternative RNA extraction method for on-site HCV detection. Boiling with DEPC-treated distilled water was not appropriate for low HCV load samples and boiling with a lysis buffer was not recommended.

## Introduction

Hepatitis C virus (HCV) is a leading cause of chronic liver disease, cirrhosis, and liver cancer. The current oral direct-acting antiviral (DAA) combinations can cure 95-99% patients [1], but the majority of HCV-infected people are unaware of their infection status. Testing for antibody to HCV (anti-HCV) is insufficient to diagnose a current HCV infection or ongoing HCV replication. HCV RNA testing is needed. Real time reverse transcription polymerase chain reaction (RT-PCR) is commonly used for HCV RNA detection/quantification in clinical practice. However, this test is rarely available for an on-site diagnosis, especially in remote settings since it requires specific reagents and instruments with high costs. The development of reverse transcription loop-mediated isothermal amplification assay (RT-LAMP) can facilitate the access to molecular testing of HCV. Indeed, the clinical sensitivity and specificity of RT-LAMP has been previously reported as high as 90-100% for HCV detection [2-4]. RT-LAMP also stands out in terms of rapidity, simplicity, cost-effectiveness, and accessibility, making it ideal for field or point-of-care use in remote settings where sophisticated and expensive equipment are not available. However, this technique still requires an effective nucleic acid extraction and purification steps. Currently, available commercial nucleic acid extraction kits, mostly based on a silica-based column extraction methodology, are expensive and require a centrifugation process, limiting the use of RT-LAMP for on-site diagnosis or in the field. In this study, we evaluated the efficacy of various simple RNA extraction methods from plasma samples for HCV detection using RT-LAMP and confirmed the presence of HCV RNA in extracted samples with real time RT-PCR.

## Materials and methods

This study used leftover plasma samples of 50 HCV-infected individuals who had HCV viral load testing and genotyping for clinical care at the Faculty of Associated Medical Sciences, Chiang Mai University. HCV viral load was initially measured using a commercial real-time RT-PCR assay (COBAS AmpliPrep/COBAS TaqMan HCV Test). The study was approved by the Human Experimentation Committee (Number 17/64) and the Institutional Biosafety Committee (Number CMUIBC0363004) of Research Institute for Health Sciences, Chiang Mai University, Thailand.

Viral RNA was extracted from plasma samples using four different methods: 1) NucleoSpin RNA Virus (Macherey-Nagel, Germany) and; 2) NucleoMag Virus kit (Macherey-Nagel, Germany) following the manufacturer’s recommendations; 3) Boiling with water, 100 μL plasma were mixed 1:1 with Diethyl Pyrocarbonate (DEPC)-treated distilled water (Invitrogen, USA) and boiled at 95°C for 10 min [5], the mixture was centrifuged at 8,000 xg for 1 min and supernatant was collected; 4) Boiling with lysis buffer, as above but 100 μL plasma were mixed 1:1 with a commercial lysis buffer (Lysis Buffer RAV1, Macherey-Nagel, Germany). Yield and purity of RNA extracted according the four methods were measured using the NanoDrop™ 2000/2000c Spectrophotometer.

For RNA detection by real time RT-PCR, four μL of viral nucleic acid extract were amplified with 400 nM of each HCV forward and reverse primers, and 100 nM of HCV probe [6] of the sensiFAST Probe Lo-ROX One-Step kit. Amplification was performed on the Applied Biosystems 7500 instrument as follows: 45°C for 10 min; 95°C for 2 min; 40 cycles of 95°C for 5 sec and 60°C for 20 sec. The fluorescence signal was measured at 60°C of each cycle. For RNA amplification by RT-LAMP, five μL of nucleic acid extract were used as previously described [2]. Briefly, RT-LAMP reaction was processed at 65°C for 60 min. The result was visualized with the naked eye based on the color change of reaction mixture induced by pre-added hydroxynaphthol blue. The samples that turned sky blue were considered as positive, while those that remained purple were considered as negative. To evaluate the analytical sensitivity of the four extraction methods, 10-fold serial dilutions of plasma with HCV viral load of 10^6^ IU/mL in 1X phosphate buffer saline were prepared. Aliquots of each dilution were extracted by the four extraction methods. All RNA extracts were tested for HCV RNA by both real time RT-PCR and RT-LAMP. Each reaction was performed in triplicate.

## Results

The four extraction methods showed considerably variable quantities of RNA, boiling plasma samples with a commercial lysis buffer provided the highest yield of RNA concentration 695.37 ng/μL and boiling with DEPC-treated distilled water the lowest yield, 0 ng/μL. The purity ranged from 0.14 to 3.46 and was the highest with the silica membrane-based extraction (Table 1). Analytical sensitivity on triplicate of serial dilutions of HCV RNA extracts from silica-based membrane and magnetic beads were all detected by real-time RT-PCR with similar cycle threshold (Ct) and by RT-LAMP at the 10^6^ concentrations. The numbers of triplicates detected by both techniques decreased when initial concentrations were 10^5^ or lower. RNA extracts obtained after boiling in DEPC-distilled water were less well detected at higher Ct than the silica-membrane and magnetic beads RNA extracts. Though boiling with a commercial lysis buffer gave the highest RNA yield, none of the RNA extracts gave a signal by real-time RT-PCR, independently of the initial HCV RNA concentration. The results from RT-LAMP could not be interpreted since the color turned from purple to sky blue immediately after adding the extracted RNA to the reaction mix. Thus, boiling with a commercial lysis buffer was not further evaluated.

**Table 1.**
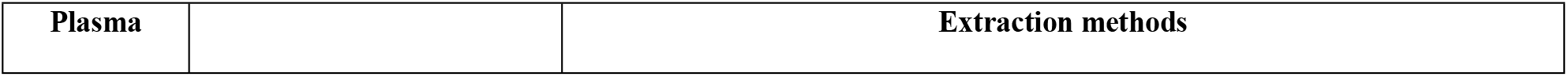
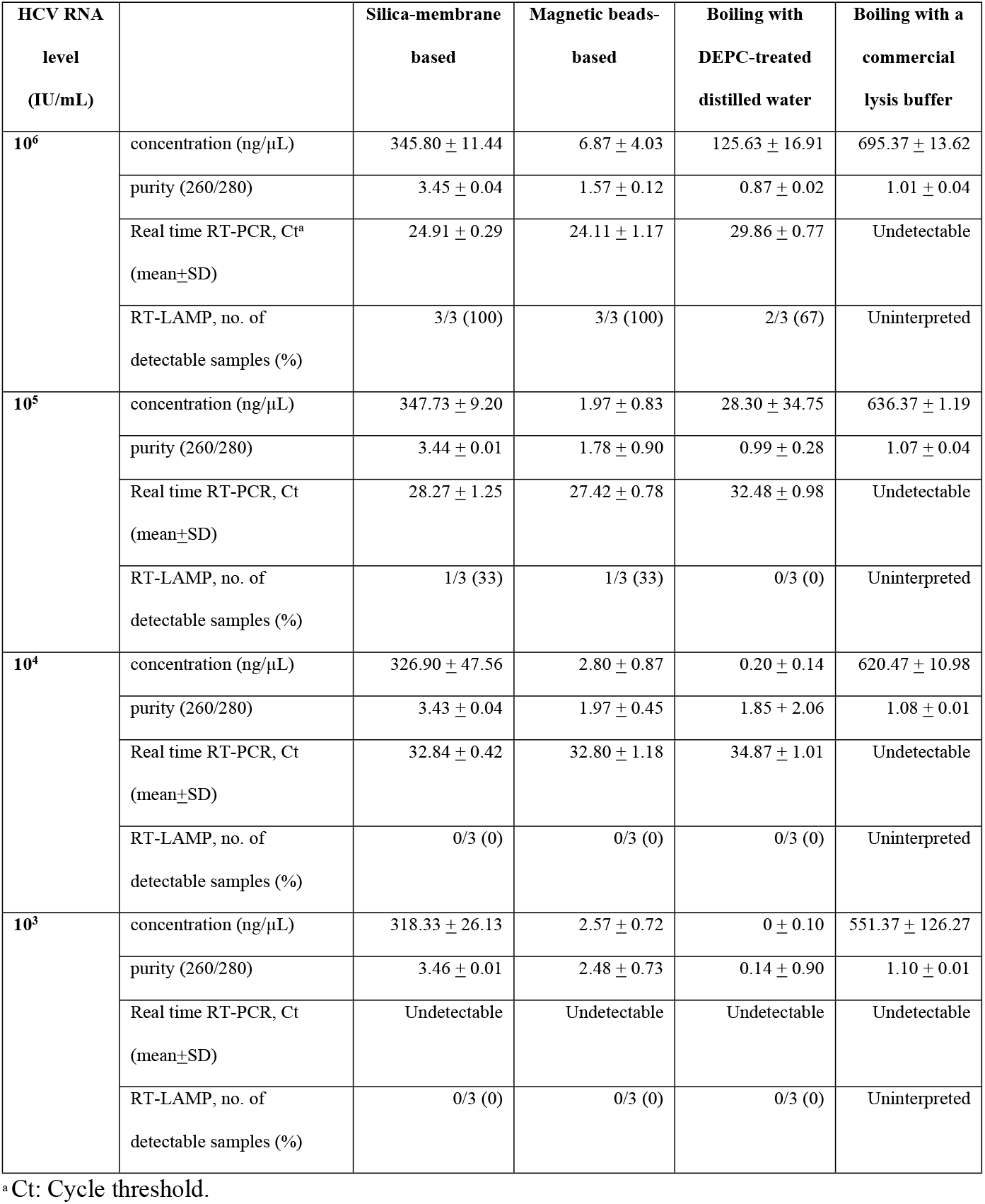
Efficacy of four different extraction methods for molecular diagnosis of HCV in plasma.

Using real time RT-PCR as measurement, RNA extracted from plasma samples using the silica-membrane-based or magnetic beads-based extraction methods gave positive scores in 50/50 (100%) (Table 2 and S1 Table). The RNA extracted from boiling with DEPC-treated distilled water gave 76% detectable rate (38/50). Using RT-LAMP for HCV detection, RNA extracted with the silica-membrane method showed the best results with 66% detectable rate (33/50), while the magnetic beads-based method showed 62% (31/50). Boiling samples with DEPC-treated distilled water provided the less number of samples with HCV RNA detected; only 7 of 50 (14%) tested positive (Table 2 and S1 Table). RT-LAMP results according to the RNA extraction methods, are shown in Fig 1. The analysis of correlation between Ct values of RNA extracted from the three different extraction methods showed an excellent correlation between Ct values of samples extracted with silica-membrane based vs magnetic beads-based methods (R^2^ = 0.88, Fig 2A). The correlation was less good between Ct values of samples extracted using silica-membrane based vs boiling with DEPC-treated distilled water methods (R^2^ = 0.60, Fig 2B).

**Table 2.** Clinical efficacy evaluation of the viral RNA extraction methods for HCV detection using real time RT-PCR and RT-LAMP.

| HCV viral load<br>(log<sub>10</sub> IU/mL) | Extraction methods |  |  |  |  |  |
| --- | --- | --- | --- | --- | --- | --- |
|  | Silica-membrane based |  | Magnetic beads-based |  | Boiling with DEPC-treated distilled water |  |
|  | Real time<br>RT-PCR, Ct<br>(mean ± SD) | RT-LAMP, no.<br>of detectable<br>samples (%) | Real time<br>RT-PCR, Ct<br>(mean ± SD) | RT-LAMP, no.<br>of detectable<br>samples (%) | Real time<br>RT-PCR, Ct<br>(mean ± SD) | RT-LAMP, no.<br>of detectable<br>samples (%) |
| 6.01-7.00 (N=10) | 24.95 ± 1.64 | 10/10 (100) | 23.19 ± 1.35 | 10/10 (100) | 29.50 ± 1.27 | 3/10 (30) |
| 5.01-6.00 (N=14) | 26.76 ± 1.13 | 13/14 (93) | 24.59 ± 0.90 | 12/14 (86) | 30.86 ± 0.93 | 4/14 (29) |
| 4.01-5.00 (N=14) | 29.89 ± 1.37 | 9/14 (64) | 27.89 ± 1.03 | 9/14 (64) | 33.83 ± 1.07 | 0/14 (0) |
| 3.01-4.00 (N=12) | 32.60 ± 1.44 | 1/12 (8) | 31.21 ± 1.23 | 0/12 (0) | 34.11 ± 0.72 | 0/12 (0) |
| Total (N=50) | 50 (100) | 33 (66) | 50 (100) | 31 (62) | 38 (76) | 7 (14) |

**Fig 1.**
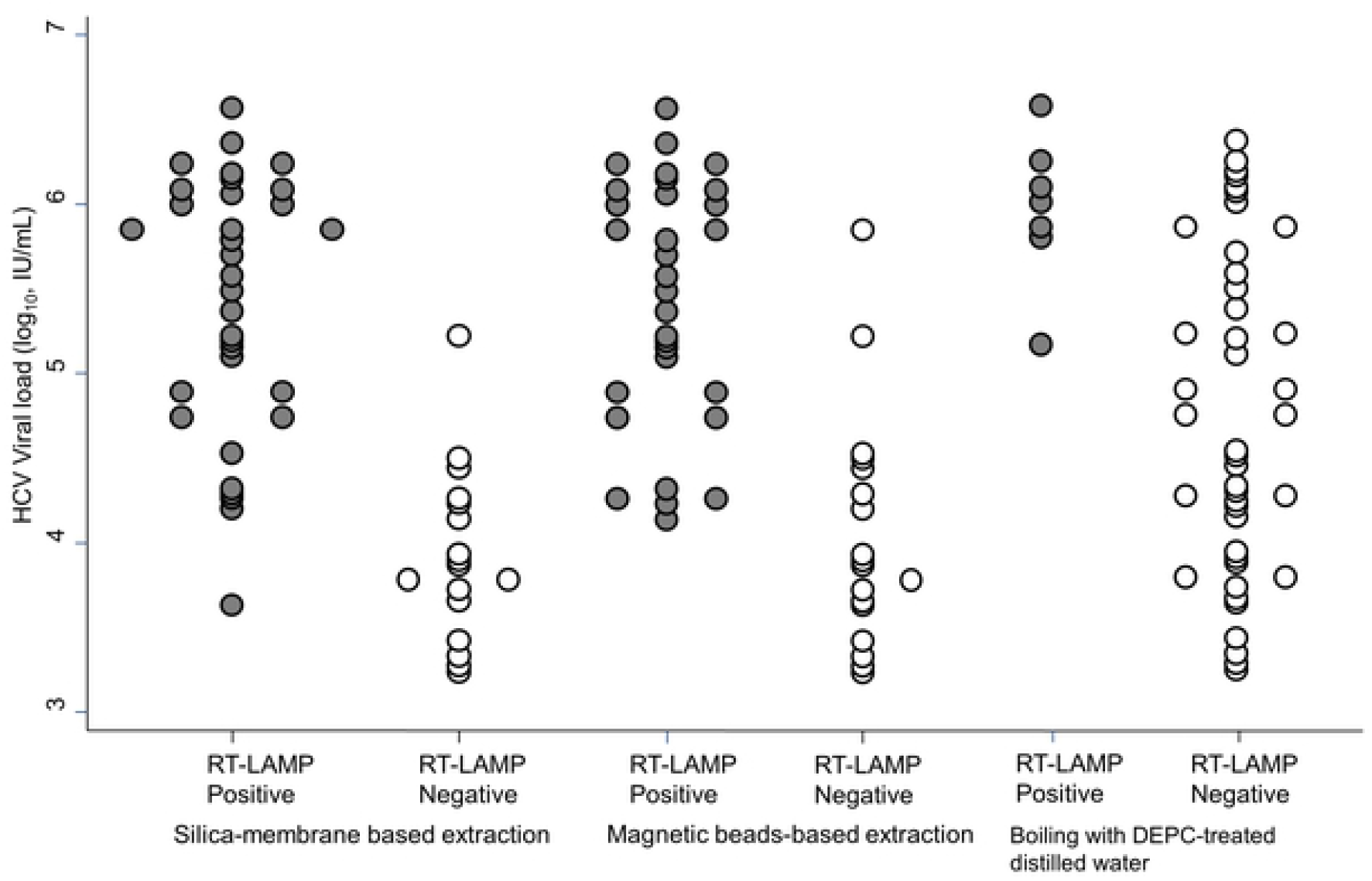
RT-LAMP results of all 50 clinical samples tested, according to the RNA extraction methods.

**Fig 2.**
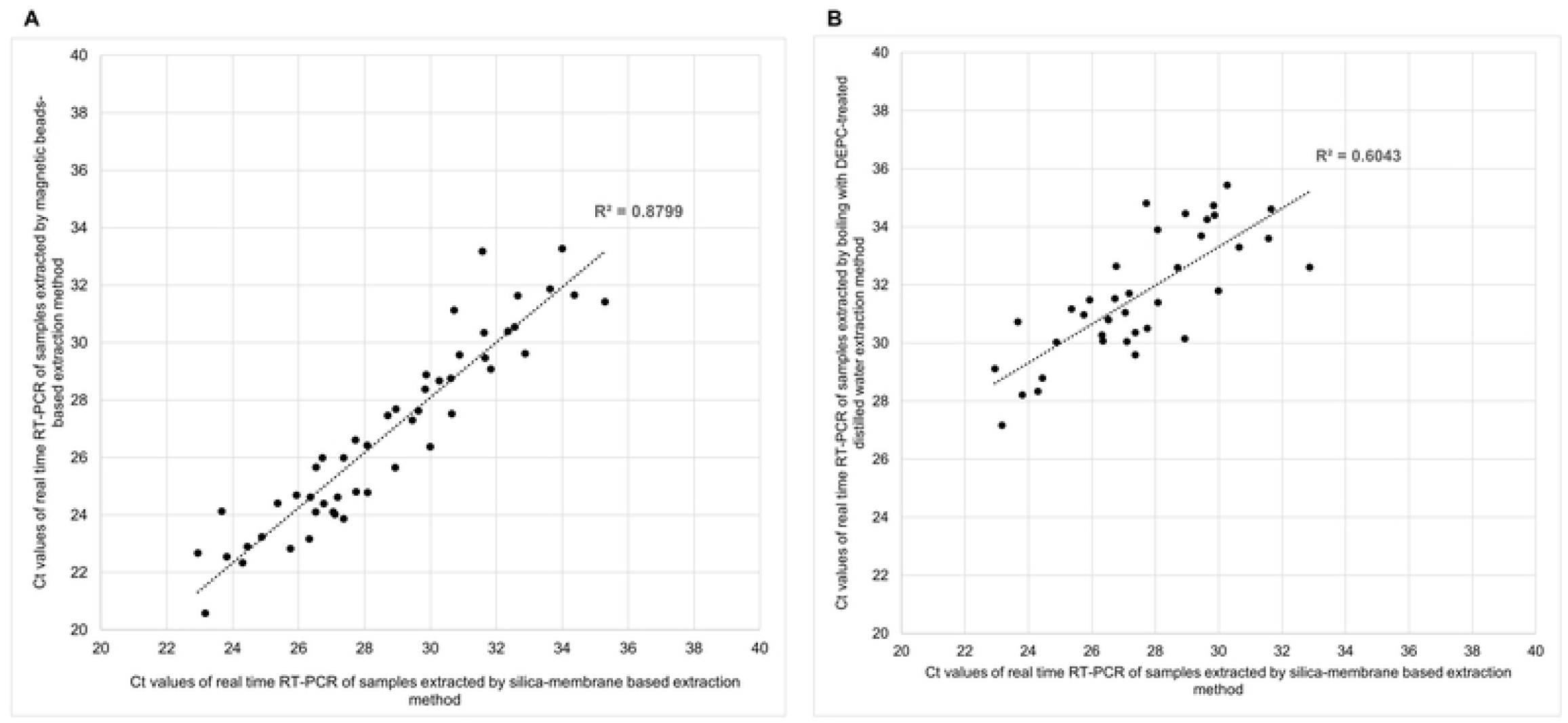
Correlation analysis between cycle threshold (Ct) values obtained by real time RT-PCR using extracted RNA (n =50) from silica-membrane based and magnetic beads-based extraction methods (A) and boiling using DEPC-treated distilled water extraction methods (B).

## Discussion

Our study showed variable sensitivities of molecular HCV detection in plasma samples depending on the RNA extraction method used. The silica-membrane based extraction method combines the selective binding properties of a silica-based membrane with the speed of microspin technology which employs a simple bind-wash-elute process. As noted, the eluates of this kit contain both viral nucleic acids and carrier RNA which may exceed the amount of authentic nucleic acids of virus when quantified by photometric method. However, the extracted RNA from this method is suitable for HCV molecular diagnosis by real time RT-PCR, and by RT-LAMP when using samples with a high viral load. Previous studies also showed that this method was suitable for RNA isolation from plasma and ready-for-use in subsequent reactions for viral detection [7, 8]. This extraction method is fast but the most expensive of the four extraction methods tested. Furthermore, it requires a centrifuge for the separation steps. The magnetic beads-based extraction method is based on the reversible adsorption of nucleic acids to paramagnetic beads. This method uses only magnetic separator for separation magnetic beads containing nucleic acids and solution. It does not need a centrifugation step but requires careful pipetting to remove the solution from the beads. In this method, 200 μL plasma sample was used, compared to only 100-150 μL of sample in the other three methods which may lead to an increase in the RNA extraction capacity. Previous studies also recommended the magnetic bead technology for viral RNA extraction from serum or plasma [9, 10]. Our results showed that Ct values from HCV samples extracted by magnetic beads-based method and silica-membrane based method correlated well. Thus, it can be used as an alternative extraction method. However, samples with low viral loads may cause a false negative result when used with RT-LAMP detection. Boiling is a simple and rapid method to release viral RNA from samples, it takes about 15 min. Previous studies suggested that simple or direct boiling without any additional purification steps can be used as an alternative RNA isolation method to detect viral infections in clinical samples [5, 11]. However, this method yielded the lowest RNA concentration and purity, as compared to others. This might be explained by a degradation of RNA during boiling and no additional step for concentration. Plasma samples showed a slightly decreased sensitivity when processed by boiling prior to amplification. When using real time RT-PCR for detection, the Ct values from HCV samples extracted by boiling were slightly higher than those of RNA extracts by silica-membrane and magnetic beads-based extraction methods and did not strongly correlate. Although the boiling method could be used as a cost-effective alternative to expensive extraction methods, it can only be used when samples have a high viral load. Boiling samples with a commercial lysis buffer provided the highest RNA concentration, a quite low purity of RNA. This might be due to the RNA carrier contained in the lysis buffer and the absence of additional step to remove the lysis buffer components or purification prior to amplification. We were unable to detect any HCV RNA in all RNA extracts and the results of RT-LAMP could not be interpreted. This may be due to an effect of inhibitors created or released by boiling. Thus, boiling plasma samples with a commercial lysis buffer cannot be used for RNA extraction in viral detection. The limitation of this study may be the relatively low number of samples and the diversity of HCV genotypes tested.

In summary, the magnetic beads-based extraction method can be used as an alternative method of plasma RNA extraction for HCV detection. This method is simple, rapid, inexpensive, and does not require a centrifugation process which makes it suitable for on-site diagnosis or in the field when combined with RT-LAMP technique. This approach will contribute to identify new HCV viremic cases and to reaching the long-term goal of HCV eradication.

## Acknowledgements

We would like to thank the Division of Clinical Microbiology, Faculty of Associated Medical Sciences, Chiang Mai University for specimen processing.

## Author contributions

**Conceptualization:** Sayamon Hongjaisee, Woottichai Khamduang

**Project administration:** Sayamon Hongjaisee

**Investigation:** Sayamon Hongjaisee, Woottichai Khamduang

**Methodology:** Yosita Jabjainai, Suthasinee Sakset

**Formal analysis:** Sayamon Hongjaisee, Yosita Jabjainai, Suthasinee Sakset

**Data curation:** Sayamon Hongjaisee

**Validation and Visualization:** Sayamon Hongjaisee, Kanya Preechasuth, Woottichai Khamduang

**Supervision:** Nicole Ngo-Giang-Huong, Woottichai Khamduang

**Writing – original draft:** Sayamon Hongjaisee, Woottichai Khamduang

**Writing – review & editing:** Sayamon Hongjaisee, Yosita Jabjainai, Suthasinee Sakset, Kanya Preechasuth, Nicole Ngo-Giang-Huong, Woottichai Khamduang

## Supporting information

**S1 Table. Clinical efficacy evaluation of the viral RNA extraction methods for HCV detection using real time RT-PCR and RT-LAMP by samples**

